# Analytical tractability of cross-scale mutualistic networks unifies persistence, collapse, and invasibility

**DOI:** 10.64898/2026.03.25.714068

**Authors:** Fernanda S. Valdovinos

## Abstract

Cross-scale integration remains a persistent challenge in ecology. Mechanistic network models have advanced this integration by linking individual behavior to community dynamics. Their complexity, however, often limits exploration to numerical simulations, which tend to be insufficient for fully unveiling the fundamental rules governing system behavior. Extracting these rules requires moving beyond numerical observation to establish exact, analytical constraints. Here, a complete mathematical analysis of a mechanistically detailed plant–pollinator model is presented. This cross-scale analysis decouples transient and equilibrium dynamics, proving that pollination strictly gates plant persistence while recruitment competition caps equilibrium abundance. The precise behavioral mechanisms scaling up to determine network stability are determined: nestedness stabilizes communities by generating floral reward gradients that guide adaptive foraging, whereas connectance destabilizes by eroding these rescue pathways. Additionally, native community persistence and biological invasions are conceptually unified; a single, multi-scale reward threshold (*R*\*) is shown to govern both native survival and alien establishment. These analytical derivations are distilled into conceptual frameworks and visual summaries accessible for empiricists interested in theory and conceptual unification. By translating numerical observations into rigorous, trait-grounded proofs, this analysis demonstrates that complex, cross-scale networks are tractable, revealing the precise conditions under which communities assemble, persist, and collapse.

**Central Question:** Can mechanistic cross-scale ecological models be made analytically tractable, and what does that tractability reveal about persistence, collapse, and invasion that simulations alone cannot?

**Answer:** A complete analytical treatment of a consumer-resource plant–pollinator model with adaptive foraging shows that cross-scale complexity can be mathematically tractable. The analysis reduces individual foraging behavior, floral reward dynamics, plant recruitment, and network topology to exact thresholds governing persistence, extinction cascades, and invasibility. These thresholds show that pollination gates plant persistence while competition caps equilibrium abundance, explain previous simulation results from first principles, and unify native survival and alien establishment under shared mathematical conditions.

**Field Map Coordinates:** Persistence and fragility; Scaling and levels; Inference, prediction, and models; Organization and emergence; Principles and theory;

## Introduction

Cross-scale integration remains one of ecology’s most persistent challenges (Levin, 1992; Peters et al., 2007). This difficulty is increasingly consequential as ecological prediction requires extrapolation beyond historical conditions. Predicting responses to novel environmental conditions requires models that connect individual behavior, physiology, and resource use to population, community, and ecosystem dynamics (Sutherland et al., 2013; Heffernan et al., 2014). Traditional single-scale, phenomenological models struggle in this role: because they rely on static parameters, they can describe systems under stable historical conditions but often fail under unprecedented perturbations such as climate warming or habitat loss (Cuddington et al., 2013; Urban et al., 2016). Mechanistic cross-scale models offer a powerful alternative by explicitly connecting lower-level processes, such as individual behavior, physiology, or resource use, to higher-level outcomes, such as population growth, community persistence, network stability, or ecosystem function. By tracing how environmental stressors alter those lower-level processes and propagating their consequences upward, such models can predict emergent community dynamics without historical precedent. Yet this mechanistic realism often comes at the cost of apparent analytical intractability: by coupling processes across biological scales, these models are typically explored through simulation, leaving unclear whether they can reveal general mathematical principles. This tension raises the central question of this paper: Can mechanistic cross-scale ecological models be made analytically tractable, and can that tractability reveal principles of persistence, collapse, and invasion that simulations alone cannot?

This tension reflects a broader debate in theoretical ecology over the relationship between model simplicity, mechanistic realism, and generality (Levins, 1966; Evans et al., 2013). Simple models have been indispensable for deriving foundational ecological principles (May, 1972; MacArthur, 1970; Allesina and Tang, 2012; Marquet et al., 2014). Yet the assumption that mechanistic complexity necessarily prevents analytical insight can be misleading. Initial complexity need not imply permanent intractability: the same cross-scale couplings that make a model difficult to analyze may also imply constraints that, once derived, make its dynamics decomposable. Here, I test this possibility using a mechanistic plant–pollinator model that links adaptive foraging, floral reward dynamics, plant recruitment, pollinator population dynamics, and network topology.

When theoretical ecology has embraced cross-scale complexity, it has uncovered mechanisms that govern emergent community dynamics. In food-web theory, models of adaptive foraging show how individual behavioral flexibility can scale up to stabilize populations, promote coexistence, and reshape the classic complexity–stability relationship (Abrams, 1992; Křivan, 1996; Kondoh, 2003; Uchida et al., 2007; Valdovinos et al., 2010). Trait-based approaches similarly show that embedding individual-level physiological constraints, such as body-mass allometry or thermal performance, can explain interaction-strength distributions, network persistence, and biodiversity responses to environmental change that simpler models miss (Brose et al., 2006; Binzer et al., 2016; Gauzens et al., 2020). These advances demonstrate the value of mechanistic cross-scale modeling, but they also highlight a recurring limitation: such models are often explored primarily through numerical simulation.

Simulations are powerful tools for discovering emergent patterns, but they can leave the mathematical machinery that generates those patterns partly opaque. Mathematical analysis can translate these numerical observations into mechanistic understanding by revealing the exact parametric boundaries, thresholds, and constraints that govern system behavior. Here, I demonstrate this point through a complete mathematical analysis of a mechanistically detailed consumer-resource model of plant–pollinator networks developed over the past decade (Box 1). The analysis reduces individual foraging behavior, floral reward dynamics, plant recruitment, pollinator population dynamics, and network topology to a small set of exact thresholds that determine whether species persist, collapse, or invade.

The resulting answer is both methodological and biological. Methodologically, the analysis shows that a complex cross-scale network model can be made analytically tractable by deriving the constraints that link behavior, rewards, visitation, and demography. Biologically, it reveals mechanisms that were hidden in simulations: plant persistence is governed by coupled pollination and recruitment thresholds, adaptive foraging reshapes the reward environment that sustains vulnerable pollinators, and network structure determines when these rescue pathways persist or fail. The same thresholds also explain why native persistence and alien establishment are governed by common mathematical conditions. Thus, analytical tractability does more than simplify the model: it exposes the biological rules by which cross-scale feedbacks organize persistence, collapse, and invasibility.

The analysis follows the cross-scale architecture of the system, deriving plant and pollinator persistence thresholds, reward dynamics, visitation balance, and network-level consequences in sequence. Because each subsection derives quantities required by the next, the equilibrium closes only when the community reward budget is reconciled with plant visitation requirements. These analytical results are then extended to transient dynamics, invasibility, and the destabilizing effects of high connectance, and are mapped directly onto previous simulation outcomes summarized in Box 2. The model structure, cross-scale feedbacks, simulation results, and new mathematical findings are summarized conceptually in Box 1, Figure 1, Box 2, and Box 3.

**Figure 1:**
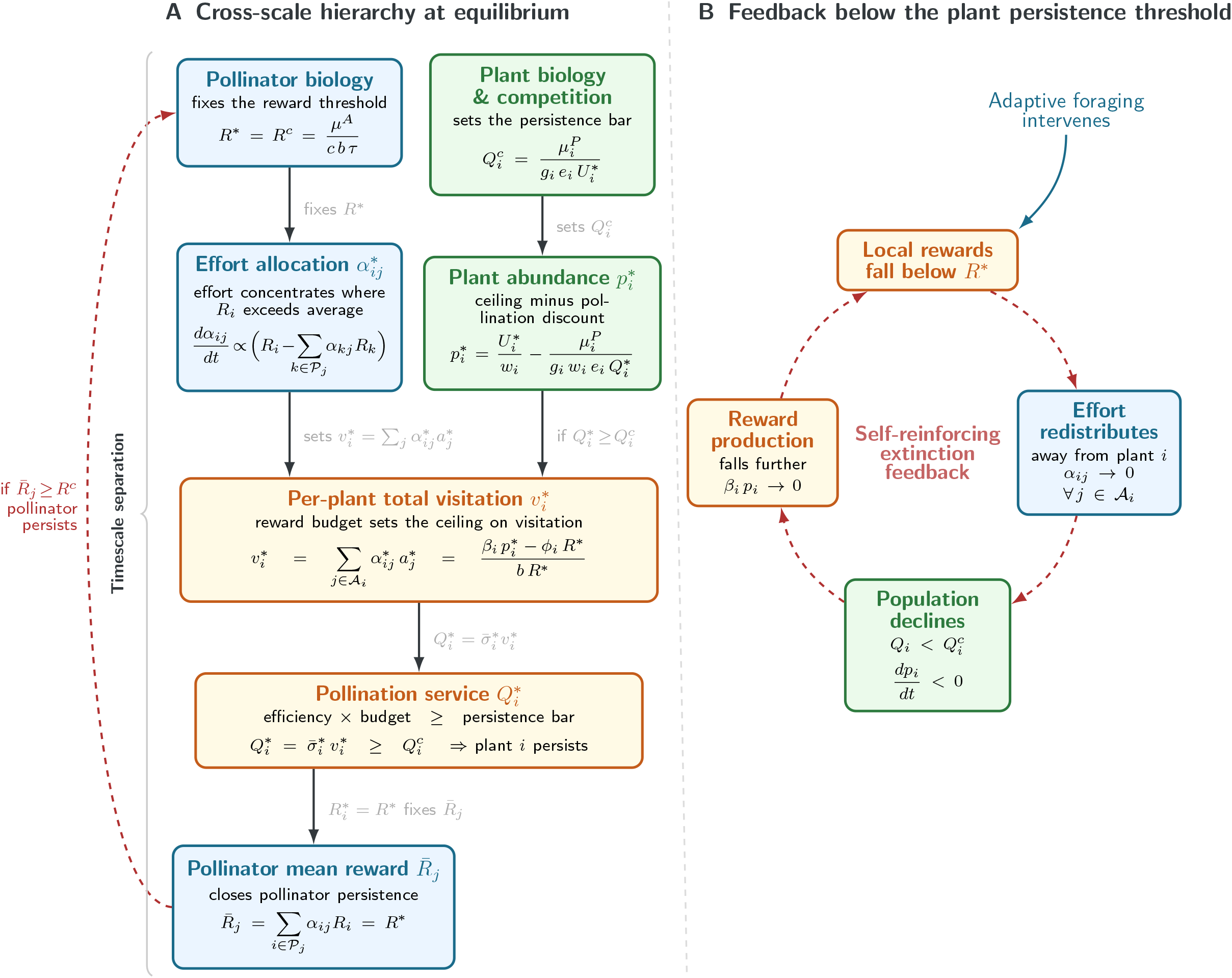
Cross-scale feedbacks determine persistence. (**A**) The mathematical hierarchy: fast behavioral adjustments in foraging allocation drive reward dynamics, which aggregate to determine slow demographic outcomes for both plants and pollinators. Pollinator biology fixes the reward threshold *R*^*c*^, and adaptive foraging equalizes standing rewards at *R*\* = *R*^*c*^, setting the level at which the community operates. Plant persistence requires the pollination service 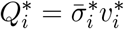 to exceed the independently set plant threshold 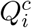. (**B**) The self-reinforcing extinction feedback: when pollination service falls below 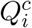, plant abundance declines, reducing reward production and causing local rewards to fall below *R*\*. Adaptive foragers then abandon the plant (*α*_*ij*_ → 0), reward production collapses (*β*_*i*_*p*_*i*_ → 0), and the plant enters irreversible decline (*dp*_*i*_*/dt <* 0). Box colors denote the biological domain: blue for pollinator traits and behavior, green for plant demography and competition, and orange for the mutualistic interface, rewards and pollination service, that couples them. See Table 1 for definitions of the cross-scale quantities.

**Table 1:** Cross-scale quantities and persistence conditions. Five derived quantities link pollinator biology, plant demography, reward dynamics, foraging behavior, and network structure. Together, they close the equilibrium and determine which species persist. Full state variables, functions, parameters, and dimensions are listed in Table **??**.

| Quantity | Formula | Biological levels | Cross-scale interpretation |
| --- | --- | --- | --- |
| Equilibrium reward $R^*$ | $R^* = \frac{\mu^A}{cb\tau}$ | Pollinator biology ($\mu^A, c, \tau$); governs $R_i$ and $\alpha_{ij}$ | Minimum reward every pollinator species needs to persist; fixed by pollinator biology alone and identical across plant species under adaptive foraging. |
| Per-plant total equilibrium visitation $v_i^*$ | $v_i^* = \frac{\beta_i p_i^* - \phi_i R^*}{b R^*}$ | Plant reward budget ($\beta_i, p_i^*, \phi_i$); pollinator biology ($R^*$); pollinator behavior and abundance ($\sum_j \alpha_{ij}^* a_j^*$) | Total effort-weighted visitation plant $i$ sustains at equilibrium; links plant abundance and reward dynamics to behavioral-level visitation. |
| Pollination threshold $Q_i^c$ | $Q_i^c = \frac{\mu_i^P}{g_i e_i U_i^*}$ | Plant vital rates ($\mu_i^P, g_i, e_i$); available recruitment niche ($U_i^*$) | Minimum pollination service plant $i$ needs to persist; set by plant biology and competitive context, but met or missed through pollinator behavior. |
| Plant equilibrium abundance $p_i^*$ | $p_i^* = \frac{U_i^*}{w_i} - \frac{\mu_i^P}{g_i w_i e_i Q_i^*}$ | Recruitment competition ($U_i^*, w_i$); plant vital rates ($\mu_i^P, g_i, e_i$); pollination service ($Q_i^*$) | Plant equilibrium separates into a competitive ceiling $U_i^*/w_i$ and a pollination-dependent mortality discount; competition caps abundance while pollination governs whether the plant persists. |
| Pollinator effort-weighted mean reward $\bar{R}_j$ | $\bar{R}_j = \sum_{i \in \mathcal{P}_j} \alpha_{ij} R_i$ | Foraging effort ($\alpha_{ij}$); reward dynamics ($R_i$); network topology ($\mathcal{P}_j$) | Mean reward pollinator $j$ experiences across its diet; whether it equals or exceeds $R^*$ determines pollinator $j$'s persistence. |

*Note.* $R_i$ is the standing reward on plant $i$ ; $R^*$ is the common equilibrium reward under adaptive foraging; $\bar{R}_j$ is the effort-weighted mean reward experienced by pollinator $j$ ; $\mathcal{R}_j = \sum_{i \in \mathcal{P}_j} R_i$ is the total reward available across pollinator $j$ 's diet; and $\mathcal{R}_j^* = R^* d_j$ is the total reward budget required across that diet when rewards are equalized. Plant species $i$ persists iff $Q_i \geq Q_i^c$ ; pollinator species $j$ persists iff $\bar{R}_j \geq R^*$ .

### Box 1

**A consumer-resource model of plant–pollinator networks**

This work analyzes the consumer-resource model of Valdovinos et al. (2013), in which nodes represent species and links indicate which pollinator *j* visits which plant *i*. The model tracks four coupled quantities: plant abundance *p*_*i*_, pollinator abundance *a*_*j*_, floral rewards *R*_*i*_, and the per-capita foraging effort *α*_*ij*_ that pollinator *j* allocates to plant *i*:

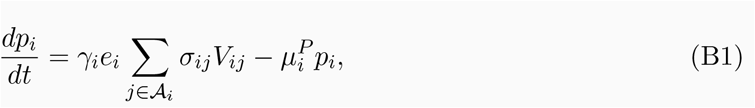

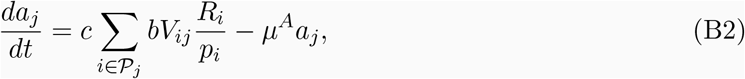

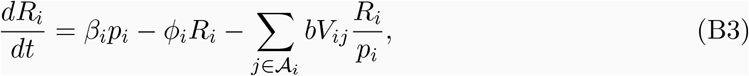

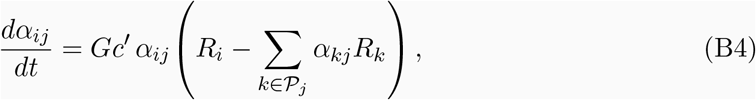

with 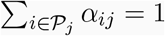 over all *j*’s plants (*P*_*j*_), and visitation rate *V*_*ij*_ = *α*_*ij*_ *τ a*_*j*_ *p*_*i*_, where *τ* = 1. See Appendix Table A1 for model notation, parameter definitions, and dimensions.

**Plant dynamics** (Eq. B1). Recruitment passes through three filters: visit quality 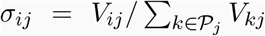, seed-set fraction *e*_*i*_, and germination–competition fraction 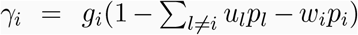, where *u*_*l*_ and *w*_*i*_ are inter- and intraspecific competition coefficients for non-pollinator resources. Mortality occurs at per-capita rate 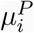. *A*_*i*_ is *i*’s pollinators set.

**Pollinator dynamics** (Eq. B2). Recruitment is proportional to rewards extracted per visit (*bV*_*ij*_*R*_*i*_*/p*_*i*_), converted to adults at rate *c*, with mortality rate *µ*^*A*^. All pollinator species share the same parameter values; differences in abundance arise solely from network structure.

**Rewards dynamics** (Eq. B3). Rewards accumulate through plant production (*β*_*i*_*p*_*i*_), saturated by self-limitation (*ϕ*_*i*_*R*_*i*_), and are depleted by pollinator consumption (*bV*_*ij*_*R*_*i*_*/p*_*i*_).

**Foraging effort dynamics** (Eq. B4). Effort shifts toward plants offering above-average rewards and away from those offering below-average rewards, at rate *Gc*^*′*^ (*c*^*′*^ = *c* · *b* · *τ*), until rewards equalize across all visited plants.

### Box 2

**Key results from simulation studies**

Results obtained via extensive simulations of the model (Box 1). The present mathematical analysis proves and mechanistically explains these results. In parenthesis is the mapping to the novel findings derived from first principles in Box 3 that explains each of them.

**R1. Adaptive foraging drives niche partitioning across scales**. When pollinators forage adaptively, foraging effort shifts toward specialist plants, which retain more unconsumed floral rewards than heavily visited generalists. This redistribution produces niche partitioning of floral rewards among pollinators and of pollination services among plants, enabling vulnerable specialist species to persist (Valdovinos et al., 2013, 2016; Valdovinos and Marsland III, 2021). *(Findings 2, 3, 5)*

**R2. Adaptive foraging reverses the effects of network structure**. Under fixed foraging, nestedness destabilizes and connectance stabilizes plant–pollinator communities. Under adaptive foraging, both effects reverse: nestedness becomes stabilizing and connectance destabilizing. The same network structure produces opposite outcomes depending on foraging behavior (Valdovinos et al., 2016). *(Finding 5)*

**R3. Species traits determine the invasion success of both pollinators and plants**. Across extensive simulations, invader traits consistently override network properties in predicting establishment. For new pollinators, high foraging efficiency is the strongest predictor of success (Valdovinos et al., 2018). For new plants, sufficiently high rewards production (*β*) secures them enough high-quality visits to invade (Valdovinos et al., 2023). In both cases, network properties predict the subsequent impacts on native species *(Findings 2 and 4)*

**R4. Pollination governs plant transient persistence but not final abundance**. Simulations revealed a counterintuitive result: plant final abundance is driven entirely by recruitment competition, thus pollination seemed inconsequential. However, pollination acts as a strict gatekeeper during the transient phase. A plant population must secure sufficient high-quality visits before its rewards drop below *R*\*. If rewards fall below this threshold, pollinators abruptly abandon the plant, triggering irreversible extinction. Thus, transient dynamics ultimately dictate community composition and plant invasion success (Valdovinos and Marsland III, 2021; Valdovinos et al., 2023). *(Findings 1, 2, and 4)*

**R5. Nestedness, moderate connectance, and adaptive foraging jointly maximize pollination services**. Empirical network structures (nested and moderately connected) produced the highest pollen deposition rates (here, pollination services, 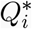) when paired with adaptive foraging. Increasing connectance beyond empirical levels reduced pollination services, whereas adaptive foraging consistently enhanced them relative to fixed foraging across all network structures. Maximizing these services depended on the interaction between network structure and behavior (Valdovinos, 2025). *(Finding 5)*

### Model Framework and Analytical Results

The analysis presented defines two foraging scenarios to bracket the behavioral range of the model (Box 1, Eq. B4). In the fixed foraging baseline, each pollinator *j* allocates effort uniformly across its diet, such that *α*_*ij*_ = 1*/d*_*j*_ for all *i* ∈ *P*_*j*_, where *d*_*j*_ is the pollinator’s degree (i.e., number of plant species it visits) and *P*_*j*_ is the set of all plant species *j* visits. That is, network topology alone (or network structure in ecological literature) determines how visits are distributed. In the adaptive foraging scenario, foraging efforts change according to Eq. B4: pollinators continuously redirect effort toward plants offering above-average rewards until rewards equalize at a common level *R*\*. Fixed foraging serves as a structural null model, but the core of the analysis focuses on the adaptive scenario to reveal how cross-scale feedbacks determine community persistence. To anchor this analysis, Table 2 provides a comprehensive side-by-side comparison of the emergent outcomes under both behavioral regimes.

**Table 2:** Equilibrium outcomes with and without adaptive foraging. *R*\* denotes the common reward threshold, 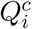 and 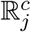 the plant and pollinator persistence thresholds, and *d*_*j*_ pollinator degree.

| Feature | Without adaptive foraging | With adaptive foraging |
| --- | --- | --- |
| *Core Mechanisms* |  |  |
| Reward distribution | Unequal across plants; determined by network topology | Equalized at $R^*$ across all visited plants |
| Foraging effort allocation | Fixed at $\alpha_{ij} = 1/d_j$ | Redistributed toward above-average rewards until $R^*$ is reached |
| *Species Persistence* |  |  |
| Specialist plants | Fall below $Q_i^c$ due to insufficient visitation | Rescued by pollinator effort shifting to their less-consumed rewards |
| Generalist plants | Receive high visitation from most pollinators, keeping $Q_i^*$ well above $Q_i^c$ | Abandoned by pollinators chasing higher rewards; survive only via specialists with no alternative links |
| Specialist pollinators | Fall below $\mathbb{R}_j^c$; rewards on plants shared with generalists are depleted by over-visitation | Rescued indirectly; generalists vacate shared plants, freeing rewards for specialists |
| Generalist pollinators | Effort fixed by topology; broad diet breadth keeps $\mathbb{R}_j^*$ above $\mathbb{R}_j^c$ despite unequal rewards | Meet $\mathbb{R}_j^c$ efficiently, driving the redistribution that rescues specialist species |
| *Network-Level Effects* |  |  |
| Degree-scaled threshold | $\mathbb{R}_j^* = R^* d_j$ (unmet for specialists) | $\mathbb{R}_j^* = R^* d_j$ (met efficiently) |
| Nestedness effect | Destabilizes specialists; concentrates visitation on generalists | Stabilizes generalist plants; niche partitioning emerges from behavior |
| Connectance effect | Stabilizes specialists by increasing both visitation and reward access | Destabilizes generalist plants by giving specialist pollinators alternative links |
| Extinction mechanism | Misallocation of fixed effort | Self-reinforcing cascade once $R^*$ is crossed (Box 3, Fig. 1B) |

#### 1. Plant equilibrium density, persistence threshold, and the roles of pollination and competition

Dividing Eq. B1 by *p*_*i*_, the plant per-capita growth takes the compact form

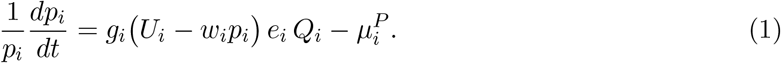

where 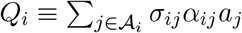 is the quality-weighted pollination service, and *U*_*i*_ ≡ 1 − ∑_*l* ≠*i*_ *u*_*l*_*p*_*l*_ is the fraction of the recruitment niche remaining after interspecific competition. Two equilibria derive from Eq. (1): the trivial 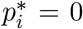, which represents local extinction, and a non-trivial equilibrium 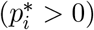 satisfying 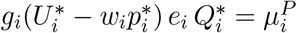, giving

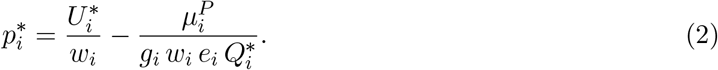

The non-trivial equilibrium separates cleanly into two terms: the first, 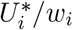, is the competitive ceiling set by the interspecific and intraspecific plant recruitment environment; the second is a mortality discount that shrinks as pollination service 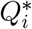 improves (Table 1). Equilibrium abundance therefore increases with 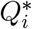 but saturates toward 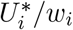 as 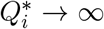, revealing the distinct life-cycle roles of each process: pollination gates seed production and therefore governs whether a plant can persist and recruit at all, while competition for establishment determines the ceiling on adult abundance that no amount of additional pollination can exceed.

Imposing 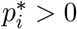 on Eq. (2) yields a minimum pollination threshold below which plant *i* cannot persist:

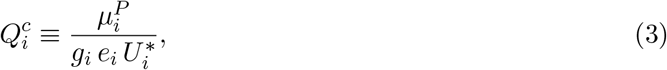

such that plant *i* persists if and only if 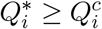. The threshold 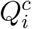 quantifies how much pollination a plant needs to persist: it rises with increased mortality and interspecific competition pressure (low 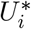), and falls with more efficient germination and seed (Table 1). Notably, intraspecific competition *w*_*i*_ is absent from 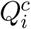: self-crowding is negligible when the plant is rare, and thus plays no role in invasion or recovery from low density.

#### 2. Pollinator persistence threshold and equilibrium density

Dividing Eq. B2 by *a*_*j*_, the pollinator per-capita growth takes the compact form

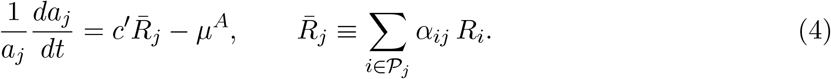

where *c*^*′*^ ≡ *cbτ* is the effective conversion rate from floral rewards to pollinator growth and 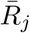 is the foraging-effort-weighted mean reward available to pollinator *j*. Imposing 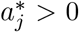 on Eq. (4) yields a minimum effort-weighted reward threshold below which pollinator *j* cannot persist:

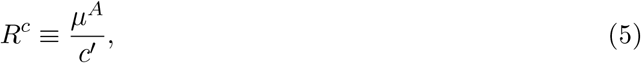

with pollinator *j* persisting if and only if 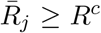. The threshold *R*^*c*^ quantifies how much effort-weighted floral reward a pollinator needs to persist: it rises with increased mortality and falls with more efficient reward-to-animal conversion (Table 1). Notably, *R*^*c*^ is the same for every pollinator species because all share the same parameters *µ*^*A*^ and *c*^*′*^. Given these shared parameters, differences in persistence and abundance therefore arise from differences in 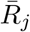, which is determined by network structure (the plant species in *j*’s diet, *P*_*j*_) and how foraging effort (*α*_*ij*_) is distributed across those plants (see next subsection).

The reward dynamics (Eq. B3) determine pollinator equilibrium abundance, which cannot be obtained from the pollinator per-capita equation (Eq. 4), because it contains no self-limiting term. Setting *dR*_*i*_*/dt* = 0 and re-arranging terms gives

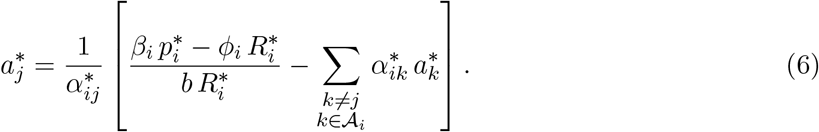

where the fraction inside brackets is the total effort-weighted visitation plant *i* can sustain, and the summation subtracts the visitation already supplied by other pollinators. The bracketed quantity is therefore the contribution that pollinator *j* must provide to plant *i*; dividing by 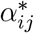 gives the abundance of pollinator *j* required to generate that contribution. The expression is not closed: 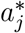 depends on the abundances of all other pollinators sharing plant *i*, and on the reward level 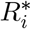 that is itself shaped by total visitation. Closure requires the reward dynamics, developed in the next subsection.

#### 3. Rewards and foraging: *R*\* as pollinator persistence threshold, community budget, and the cross-scale mediator of specialist pollinator fate

Under the replicator dynamics (Eq. B4), effort shifts continuously toward plants offering above-average rewards and away from those offering below-average rewards. This redistribution continues until rewards across all plants reach the same value, which determines the equilibrium for both rewards and foraging efforts. Because the weights *α*_*ij*_ sum to one over *P*_*j*_, the term 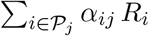 is simply the foraging-effort-weighted mean reward available to pollinator *j*. When all plant rewards equalize to a common value *R*\*, this weighted mean collapses to *R*\* regardless of diet breadth or effort distribution:

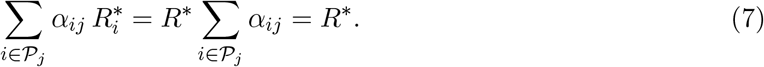

Pollinator persistence requires 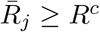 (Eq. 5), but any persisting pollinator at equilibrium must have zero net growth, so this condition holds with equality. Substituting the equalized reward level *R*\* into Eq. (5) therefore gives

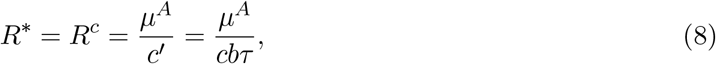

which holds simultaneously for every persisting pollinator. Eq. (8) defines both a population-level persistence condition and a community-level constraint. At the population scale, each pollinator species must harvest *R*\* per-capita to persist. At the community scale, summing *R*\* over all plant species in *j*’ diet, reveals the minimum total reward budget *j* requires to persist (Appendix):

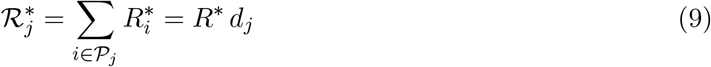

which scales linearly with *j*’s degree, *d*_*j*_ (Table 1). This degree-scaled budget is a cross-scale emergent property: pollinator biology shapes the reward landscape, and network degree sets the persistence requirement. It is also an instance of the ideal free distribution (Fretwell and Lucas, 1969), in which consumers redistribute across patches until all yield equal returns (here, *R*\*).

The threshold 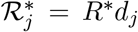 holds under both foraging scenarios (Appendix). What adaptive foraging changes is not the threshold itself, since *R*\* is fixed by pollinator biology regardless of how effort is distributed, but rather whether and how efficiently pollinators reach it. Without adaptive foraging, effort is fixed by network topology. This leaves rewards unequal across plants and visitation misallocated relative to reward budgets, a dynamic that systematically drives specialist pollinators to extinction in nested networks. Because specialists tend to visit only the most generalist plants, they face intense competition from the full pollinator community, which depletes local rewards below the persistence requirement. With adaptive foraging, generalist pollinators redirect their effort toward plants offering above-average rewards. This behavioral shift releases the visitation pressure on generalist plants, allowing their rewards to recover toward *R*\*. Consequently, the specialist pollinator’s persistence condition is met. This survival occurs not because the specialist redistributes its own effort (as a species with *d*_*j*_ = 1 has nowhere else to go), but because generalists have vacated the shared plant. Specialist persistence is therefore an indirect, cross-scale consequence of adaptive foraging, where the individual decisions of generalists determine the reward environment and ultimate survival of specialists.

#### 4. Visitation, 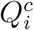, and the cross-scale determinants of plant persistence

The first subsection derived the plant persistence condition 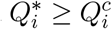, but left the actual pollination service 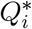 unresolved because it depends on pollinator behavior. The second and third subsections established that adaptive foraging drives the network toward a universal reward threshold (*R*\* = *R*^*c*^) dictated by pollinator persistence. This fourth subsection closes the loop by deriving the equilibrium visitation rate 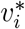 that plants receive under this uniform reward landscape. This derivation finally determines whether the plant persistence condition is satisfied.

Setting *dR*_*i*_*/dt* = 0 and substituting 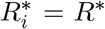 yields the equilibrium constraint on per-capita visitation (Table 1):

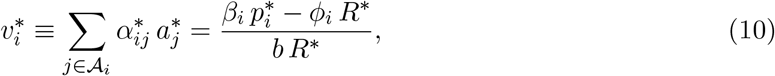

The left-hand side represents the total foraging effort directed at plant *i*, weighted by pollinator abundance. This quantity is shaped entirely by pollinator behavior and network topology. The right-hand side defines the effort-weighted pollinator abundance that the plant’s own reward budget can sustain at standing crop *R*\*. At equilibrium, these two sides must be equal. The net reward production rate 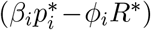 must exactly balance the reward removed per unit time 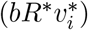. This is the mutualistic analog of Tilman’s supply-consumption balance (Valdovinos and Marsland III, 2021), expressed here as a reward zero-net-growth isocline (ZNGI) condition. The reward budget therefore acts as a hard ceiling. Visitation cannot exceed what the plant’s reward production can sustain at *R*\*, regardless of how much effort pollinators are willing to allocate.

This reward balance closes the quantity left open in subsection 1 and motivates a decomposition of pollination service into two distinct constraints. Substituting 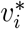 into the expression for 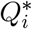 yields:

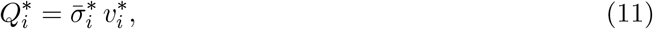

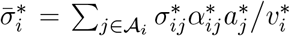 is the effort-weighted mean visit quality plant *i* receives at equilibrium. The quantity 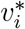 acts as a budget constraint. It sets the maximum visitation the reward budget can sustain given plant abundance and *R*\*. The quantity 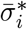 acts as an efficiency constraint. It determines how much of that visitation converts into successful pollination. This conversion depends on network topology (through 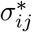) and the distribution of foraging effort. Together, Eqs. (8)–(11) fully determine the equilibrium. Pollinator biology fixes *R*\*. The reward balance fixes 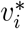. Finally, the product 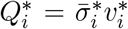 determines whether the persistence condition 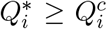 is met. Plant persistence therefore requires both constraints to be satisfied simultaneously. The reward budget must sustain sufficient visitation, and the network must convert that visitation into efficient pollination.

The consequences of this decomposition are sharpest in nested networks. Without adaptive foraging, effort is fixed by topology and concentrated on generalist plants. Specialist plants consequently receive fewer visits, and the pollinators that do visit them are generalists whose effort is spread across many species, diluting pollen quality. Both 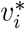 and 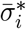 are therefore depressed. Specialist plants fall below 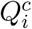 despite adequate pollinator abundance in the network, and go extinct while generalists persist. With adaptive foraging, effort continuously redistributes toward plants retaining above-average rewards. Specialist plants, less heavily visited and therefore carrying more unconsumed rewards than generalists, attract increasing foraging effort as rewards equilibrate across the network. Both 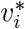 and 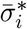 rise and the persistence condition 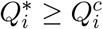 is met through the redistribution of foraging effort.

#### 5. Transient dynamics, cascades, invasibility, and the role of connectance

Subsections 1–4 characterize the network equilibrium structure by defining the persistence thresholds 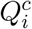 and *R*^*c*^ and establishing the universal reward level *R*\* (Eqs. 3, 5, 8). They end by decomposing pollination service into distinct budget and efficiency constraints (Eqs. 10, 11). This subsection shifts focus to the dynamics between equilibria. It examines how perturbations propagate through the network and why some state transitions are irreversible, and explores what these same persistence thresholds imply for the establishment of new species.

When a perturbation (e.g., pollinator loss) reduces *Q*_*i*_ below 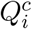 (Eq. 3), a self-reinforcing extinction feedback is initiated (Figure 1B). Declining plant abundance *p*_*i*_ reduces reward production *β*_*i*_*p*_*i*_. This drop depresses standing reward *R*_*i*_, which in turn reduces visitation further. Recovery requires crossing 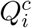 from below. However, because reward production scales with *p*_*i*_, a plant at low abundance cannot generate sufficient reward to attract pollinators and restore 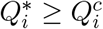. This feedback operates wherever 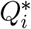 falls below 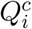: in specialist plants receiving diluted visitation from effort-spread generalist pollinators (end of last subsection), in generalist plants losing foraging effort to higher-reward alternatives under high connectance (next paragraph), and in introduced plants too sparse to generate the reward necessary for establishment (following next paragraph).

In nested networks with moderate connectance, generalist plants are the most buffered nodes against pollination failure. They receive the majority of visits in the network. More importantly, they are visited by specialist pollinators that have no alternative links and depend entirely on them. This exclusive reliance ensures a stable base of high quality visits that keeps 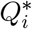 well above 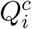. High connectance erodes this protection. When specialist pollinators acquire links to additional plant species, they redirect effort toward higher-reward alternatives. As rewards on generalist plants fall below the network mean, this behavioral shift sharply reduces visitation 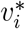 (Eq. 10). Once 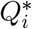 falls below 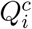, the irreversible feedback described above is triggered, converting a buffered hub into a vulnerable node.

The same thresholds that govern equilibrium persistence (Figure 1A) also define invasibility. A species establishes if and only if its per-capita growth rate is positive at low density. At low density, plant intraspecific competition is negligible. The competition term *w*_*i*_*p*_*i*_ in Eq. 1 vanishes as *p*_*i*_ → 0. Consequently, per-capita growth is determined entirely by *Q*_*i*_ relative to 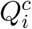. The persistence threshold and the invasibility criterion are therefore the exact same condition. Establishment faces the same self-reinforcing feedback described above (Figure 1B). However, the threshold is approached from below rather than crossed from above. Low initial abundance produces little total reward. This scarcity attracts few pollinators. The lack of visitation keeps *Q*_*i*_ below 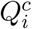, driving per-capita growth negative. Plants with sufficiently high reward production rate *β*_*i*_ can meet and exceed this threshold. Even at low abundance *p*_*i*_, a high *β*_*i*_ sustains reward levels that push *Q*_*i*_ above 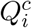 before the feedback consolidates. This dynamic selects for high reward production as a key establishment trait, independent of recruitment competitive ability or network position.

The same logic extends to pollinators. An introduced pollinator species invades if and only if the effort-weighted mean reward in the resident network exceeds *R*^*c*^ (Eq. 5). This excess reward is the strict requirement for positive per-capita growth at low pollinator density (Eq. 4). If the resident community already operates at *R*\* = *R*^*c*^ (Eq. 8), the invader finds rewards exactly at the persistence threshold. Without a surplus, the new species cannot grow. Successful invasion therefore requires one of two conditions. First, the resident community may not have fully depleted rewards to *R*\*, leaving an available surplus above *R*^*c*^. Alternatively, the invader may possess a higher reward-to-animal conversion efficiency *c*^*′*^. This higher efficiency lowers the invader’s own *R*^*c*^ below the standing reward level. Under either scenario, invasibility remains tightly bound to the equilibrium persistence threshold. This confirms that *R*\* functions simultaneously as a population-level persistence condition, a community-level reward budget constraint, and a cross-guild invasibility criterion.

##### Box 3

**Novel findings from the mathematical analysis**

The mathematical analysis presented here derives the following findings from first principles, which were unavailable to simulations or simpler frameworks.

**F1. Pollination gates plant persistence; plant competition determines equilibrium abundance**. Plant equilibrium abundance decomposes into a competitive ceiling 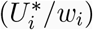 and a mortality discount that shrinks as pollination service improves (Eq. 2). Intraspecific competition *w*_*i*_ is absent from the persistence threshold 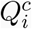: self-crowding is negligible when a plant is rare, so pollination alone gates whether a plant can persist, while plant recruitment competition determines its abundance ceiling.

**F2**. *R*\* **simultaneously functions at three biological scales**. The common reward level at which adaptive foraging equilibrates the network (Eq. 8) is at once: (i) a *population-level* persistence condition, requiring each pollinator to harvest *R*\* per-capita to avoid decline; (ii) a *community-level* reward budget, where the minimum total reward a pollinator of degree *d*_*j*_ requires scales as 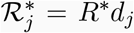 (Eq. 9); and (iii) an *invasibility criterion*, whereby a new pollinator establishes only if the reward level on visited plants exceeds *R*\* or its conversion efficiency lowers its own threshold below the resident reward level.

**F3. Pollination services decompose into quantity and quality constraints that must both be satisfied**. At equilibrium, 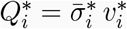 (Eqs. 10–11). Thus, plant persistence requires both constraints to be met simultaneously: the reward budget must sustain sufficient visitation quantity *and* the network must convert those visits into high-quality pollination.

**F4. The plant persistence threshold and the invasibility criterion are the same condition**. At low density, intraspecific competition is negligible (*w*_*i*_*p*_*i*_ → 0 as *p*_*i*_ → 0), leaving per-capita growth determined entirely by *Q*_*i*_ relative to 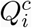 (Eq. 1). The persistence threshold and the invasibility criterion therefore collapse to a single condition: a plant establishes if and only if 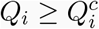. Establishment consequently faces the same self-reinforcing feedback as extinction (Figure 1B): low abundance produces little reward, which attracts few pollinators, which keeps *Q*_*i*_ below 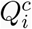 and per-capita growth negative.

**F5. Nestedness creates the topological conditions that make adaptive foraging effective; connectance erodes them**. Nestedness generates an asymmetry in unconsumed rewards between generalist and specialist plants, giving pollinators a reward gradient to follow while specialist pollinators visiting only generalist plants keep those plants above 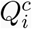. Connectance erodes both conditions simultaneously: as connectance increases, specialist pollinators acquire additional links, breaking the exclusive association that protected generalist plants, while specialist plants become accessible to a broader pollinator set, compressing the reward asymmetry that directed effort toward them.

#### 6. First principles unify structure, behavior, persistence and invasions

The analytical framework derived in the preceding subsections provides the definitive mechanistic explanation for the dynamics observed in previous simulations (Box 2). By mapping these numerical results onto the mathematical findings organized in Box 3, this analysis demonstrates that the model’s complex outcomes emerge predictably from a unified set of first principles. This subsection leverages that mapping to prove how cross-scale feedbacks dictate community-level persistence.

First, the mathematical decomposition of 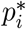 into a competitive ceiling and a pollination discount resolves a long-standing mystery from numerical simulations (Result 4 in Box 2). Without this analytical lens, pollination appeared inconsequential because it did not affect final plant abundances. The mathematics reveal a clean separation between transient and equilibrium dynamics. Pollination acts strictly as a gatekeeper during transients: it determines whether a plant persists by meeting the reward (*R*\*) and persistence 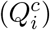 thresholds (Findings 1 and 2 in Box 3). Once persistence is secured, recruitment competition alone dictates final abundance. Thus, pollination gates whether a plant species persists, while competition caps how abundant it becomes.

Second, the derivations explicitly define how network structure and foraging behavior interact to determine persistence (Results 1, 2, and 5). The analysis reveals two distinct rescue pathways that operate in nested networks. Nestedness generates an asymmetry in unconsumed rewards, providing a gradient that drives generalist pollinators to vacate shared plants. This indirect behavioral shift allows local rewards to recover toward *R*\*, rescuing specialist pollinators from competitive exclusion (part (i) of Finding 2). Simultaneously, a direct pathway operates for plants: generalist plants receive exclusive visits by specialist pollinators, providing the high-quality visits needed to keep those plants above 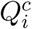. High connectance erodes both mechanisms. By providing specialist pollinators with alternative links, high connectance breaks their exclusive associations with generalist plants, triggering the irreversible extinction feedback (Figure 1B).

Finally, this framework mathematically unifies native persistence and biological invasions (Results 3 and 4). Because the invasibility criteria for both pollinators and plants are embedded in their respective persistence thresholds, invasion success is governed by the exact same transient dynamics that dictate native survival. For alien pollinators, establishment is dictated by the community-level reward budget (Finding 2): an invader succeeds if its high foraging efficiency lowers its required reward, *R*^*c*^, below the resident standing level. For alien plants, the invasibility criterion and the persistence threshold are mathematically identical (Finding 4). An invading plant succeeds by producing high initial floral rewards to rapidly attract a massive quantity of visits, which the network’s foraging behavior must then translate into sufficient quality to cross 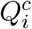.

## Discussion

The central question of this paper was whether a mechanistic cross-scale ecological model can be made analytically tractable, and whether that tractability reveals principles of persistence, collapse, and invasion that simulations alone cannot. The answer is yes. Analytical tractability is achieved for this plant–pollinator model by reducing its cross-scale feedbacks to a set of coupled thresholds linking individual foraging behavior, floral reward dynamics, plant recruitment, pollinator demography, and network topology. These constraints reveal that persistence, collapse, and invasion are not determined by topology, demography, behavior, or rewards in isolation, but by the thresholds that couple them. Community persistence is therefore not a property of any single scale. It emerges from the interaction of network topology, population dynamics, and individual behavior, and it becomes visible only when those scales are analyzed jointly.

The tractability and power of the analytical framework developed here stems from the derivation of quantities that sequentially determine others from first principles (Figure 1A). Under equal pollinator demographic parameters, pollinator biology fixes *R*\*, which sets the reward landscape and shapes foraging allocation across the entire network. The foraging dynamics that produce this equilibrium are governed by the replicator equation (Kondoh, 2003; Valdovinos et al., 2013), in which effort is continuously redistributed toward higher-reward plants until all visited species yield equal returns. Here, that individual-level process scales up: *R*\* becomes a community-level constraint that propagates through every downstream quantity in the causal chain, from the visitation each plant sustains to whether that visitation meets 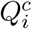, and ultimately to community persistence (Figure 1A). Because each derived quantity (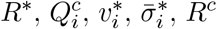; Table 1) encodes cross-scale interactions into a single analytically separable expression, the causal chain remains both biologically interpretable and mathematically modular.

The mathematical modularity of these quantities fulfills the core promise of mechanistic cross-scale modeling: explicit, parameter-grounded predictions under novel conditions (Cuddington et al., 2013; Urban et al., 2016). It isolates exactly where a given stressor enters the causal chain and predicts how it propagates across biological scales before a collapse is observed. A physiological stressor increasing pollinator mortality (*µ*^*A*^) strictly elevates the community reward threshold (*R*\*), shifting the persistence landscape in a direction and magnitude that can be calculated directly from first principles. An environmental stressor altering plant recruitment 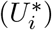 strictly elevates the pollination threshold 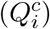, with consequences for plant persistence that are similarly traceable without simulation. This capacity for mechanism-based prediction is precisely what phenomenological models cannot provide (Cuddington et al., 2013; Evans et al., 2013) and what cross-scale analytical frameworks are uniquely positioned to deliver: tracing how a perturbation at one biological scale cascades through adaptive foraging to alter community persistence at another.

The derived thresholds also govern transient dynamics, exposing feedback processes that dominate ecologically relevant timescales (Hastings, 2004; Hastings et al., 2018). When a perturbation reduces pollination service below 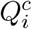, it initiates a self-reinforcing feedback: declining plant abundance reduces aggregate reward production, which depresses standing rewards and further reduces visitation. Once this feedback drives visitation sufficiently far below the plant persistence threshold, recovery requires crossing 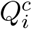 from below, which is unattainable within the closed dynamics without external rescue or parameter change. These same mathematical boundaries also govern community assembly and invasion. Whether species are being lost following a perturbation or gained during assembly, the resident reward level, *R*\*, acts as an environmental constraint against which species-specific persistence thresholds are evaluated. For pollinators, invasion requires the effort-weighted mean reward in the resident network to exceed *R*^*c*^; for plants, establishment requires pollination service to exceed 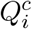 when abundance is low. Thus, *R*\* functions simultaneously as a population-level persistence condition for resident pollinators, a community-level reward budget, and an environmental filter for cross-guild invasibility. This analytical unification shows how individual-level adaptive foraging dynamics can scale up to shape community-level patterns of bio-diversity and stability. This shared threshold structure was not visible in the simulation studies that first characterized invasion dynamics in this model (Valdovinos et al., 2018, 2023), and only becomes apparent when the equilibrium structure is fully resolved analytically.

This mechanistic decomposition also clarifies how nested mutualistic networks rescue vulnerable species through complementary structural and behavioral pathways. The analytical framework presented here shows that this rescue (Valdovinos et al., 2013, 2016) is driven by two distinct mechanisms. The first is a structural pathway in which specialist pollinators, constrained to a single plant partner with *α*_*ij*_ = 1, maintain high-quality visits to generalist plants, keeping those plants above 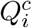. The second is a behavioral pathway in which generalist pollinators adaptively redistribute effort among co-flowering plants, thereby reshaping the reward landscape and allowing rewards on heavily contested plants to recover toward *R*\*. Together, these pathways explain why nestedness can stabilize communities: specialists provide reliable high-quality visitation to generalist plants, while adaptive foraging generates reward gradients that redistribute generalist effort. High connectance can destabilize communities by eroding these same rescue pathways, especially when additional links weaken exclusive associations and allow formerly specialized pollinators to redirect effort toward higher-reward alternatives. This result converges with independent work by Benadi and colleagues (Benadi et al., 2012, 2013), who reached similar conclusions using a structurally different model, lending cross-model support to the broader role of consumer behavior and network structure in mutualistic coexistence. Because these pathways emerge from the coupling between network topology, adaptive foraging effort, and consumer-resource dynamics, they may operate in other mutualistic systems sharing this architecture, including multiplex networks where mutualistic and antagonistic interactions co-occur (Hale et al., 2020).

Two limitations define important directions for extending this framework. First, the analysis requires parameter equality among pollinator species: conversion efficiency *c*, per-visit reward extraction *b*, and mortality rate *µ*^*A*^ are shared, producing one common *R*\* for all pollinators. This assumption is necessary here for isolating the role of network structure. Under Tilman’s *R*\* rule, allowing these parameters to vary would introduce fitness differences that drive competitive exclusion within shared-resource subsets (Tilman, 1982; Chesson, 2000). Without counteracting stabilizing mechanisms, the pollinator with the lowest *R*\* would exclude its competitors, making survival depend on parameter asymmetry rather than network architecture (Valdovinos et al., 2013, 2016; Valdovinos and Marsland III, 2021). Yet real pollinators differ substantially in body size, metabolic rate, and mortality (Heinrich, 1979; Greenleaf et al., 2007; Brown et al., 2004; Kelemen et al., 2019). Future work should therefore identify how adaptive foraging interacts with stabilizing mechanisms such as predation, environmental variability, or storage effects to allow natural networks to persist despite the *R*\* hierarchies imposed by trait variation. Second, the model assumes a fixed potential network topology. Pollinators can redistribute effort among established partners, but entirely novel interactions cannot form. In natural systems, species may sometimes avoid extinction by foraging outside their historical network or by shifting phenology (e.g., Burkle et al., 2013; CaraDonna et al., 2017; Bartomeus et al., 2011; Gaiarsa et al., 2021). Computational extensions of this (Glaum et al., 2021) and other modeling frameworks (Ramos-Jiliberto et al., 2012; Kaiser-Bunbury et al., 2010) show that such topological and temporal flexibility can buffer communities against coextinction. Incorporating these dynamic rescue mechanisms into the analytical boundaries derived here is an important next step toward predicting network resilience under severe perturbation.

The implications of this analysis should extend beyond this specific model of mutualism because the mathematical architecture developed here applies more broadly to ecological networks characterized by consumer-resource interactions and adaptive foraging. In any system where consumers dynamically reallocate effort to maximize returns, their behavioral equilibrium can act as a mathematical constraint on the underlying population dynamics. By using this behavioral constraint to decouple demographic equations, this modular approach offers a route toward deriving exact persistence boundaries for complex food webs and other mutualistic networks. The exact solutions presented here therefore serve as a proof of concept for an approach that accepts mechanistic complexity as a starting point, opens the black box of numerical simulation, and analytically recovers the biological conditions under which complex ecological networks persist, assemble, invade, and collapse. In doing so, this work shows that tractability is not merely a mathematical convenience. It is a route to biological discovery: a way to identify the thresholds, feedbacks, and cross-scale constraints that determine which ecological systems can persist and why they become fragile.

## Appendix

### A snote on the degree-scaled reward budget

For any pollinator *j*, setting 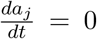 in the pollinator dynamics yields the general persistence condition

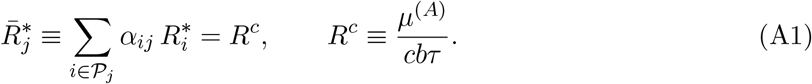

where 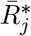 is the effort-weighted mean reward experienced by pollinator *j* at equilibrium, and *R*^*c*^ is the minimum effort-weighted reward required for pollinator persistence.

The total reward available across pollinator *j*’s diet is

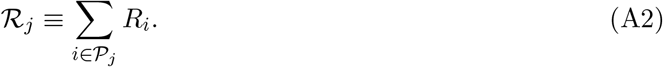

The corresponding persistence budget, 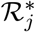, is the total reward across the diet required for 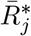 to meet the threshold *R*^*c*^.

#### Without adaptive foraging

Foraging effort is distributed uniformly, *α*_*ij*_ = 1*/d*_*j*_ for all *i* ∈ *P*_*j*_. Substituting this fixed allocation into Eq. (A1) gives

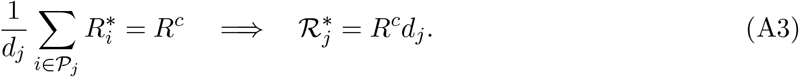

Thus, under fixed foraging, persistence requires the average reward across pollinator *j*’s diet to reach *R*^*c*^, or equivalently for the total reward available across that diet to reach *R*^*c*^*d*_*j*_.

#### With adaptive foraging

Under adaptive foraging, effort is redistributed until reward returns are equalized across visited plants. Therefore, for every plant retained in pollinator *j*’s diet at equilibrium,

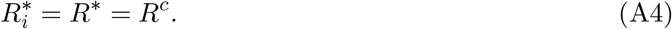

Summing over the *d*_*j*_ plants in the diet gives

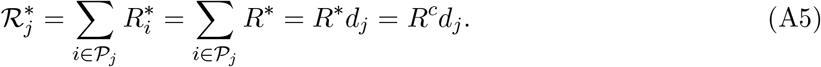

#### Interpretation

Eqs. (A3)–(A5) show that the total reward budget required for persistence scales linearly with pollinator degree under both foraging scenarios:

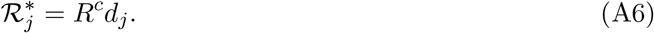

The scenarios differ, however, in how this budget is met. Without adaptive foraging, persistence depends on whether the fixed allocation exposes pollinator *j* to a sufficiently high average reward across its diet. With adaptive foraging, effort continuously redistributes until rewards equalize at 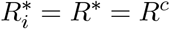 across visited plants. The consequences of this difference for specialist and generalist persistence in nested networks are developed in the main text.

**Table A1:**
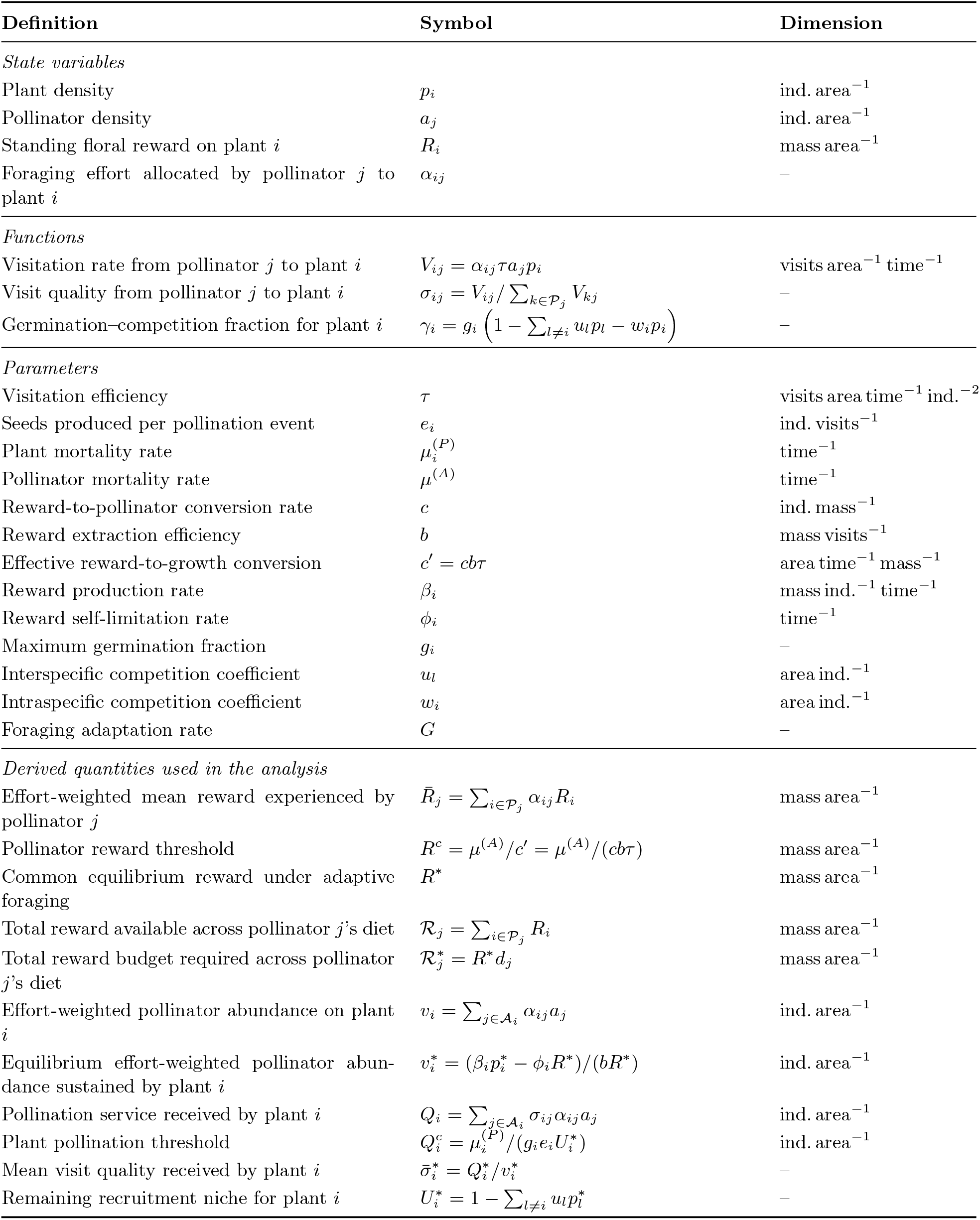
Model notation, parameters, and derived quantities. *P*_*j*_ is the set of plants visited by pollinator *j, A*_*i*_ is the set of pollinators visiting plant *i*, and *d*_*j*_ = |*P*_*j*_| is the degree of pollinator *j*.

## Notes

### Competing Interest Statement

The authors have declared no competing interest.

### Summary of Updates

This version of the manuscript better describes the question asked by the study and the answer that the study provides to such question. It also improves the introduction and the discussion.

### Funding

- U.S. National Science Foundation: DEB-2129757

